# Costs and economic impacts of expanding marine protected area systems to 30%

**DOI:** 10.1101/2022.11.20.517276

**Authors:** Anthony Waldron, Ryan Heneghan, Jeroen Steenbeek, Marta Coll, Kim J. N. Scherrer

## Abstract

International proposals for marine biodiversity seek to expand marine protected area (MPA) coverage from 8% to 30%, known as 30×30. Quadrupling MPA coverage implies considerably higher MPA system costs and governments need early knowledge of these to inform debate. Ambitious MPA expansion also implies large potential losses or “opportunity costs” for fishers, putting pressure on governments to compromise and permit some fishing inside protected areas (a mixed high/low protection system). Crafting a balanced compromise needs to be informed by model projections of future fisheries outcomes under different protection regimes, climate change scenarios and behavioural adaptations. Here, we develop the first models for management costs at national MPA-system scale. We create scenarios of 30×30 at different compromises around protection strictness. We then examine how both MPA costs and opportunity costs vary with strictness, by simultaneously applying our management cost models and two Marine Ecosystem Models. We find that a no-take (high protection) MPA system could cost just $2 billion/year for the developing world and ~$8 billion overall, but would also create opportunity costs several times larger. A compromise mix of high and medium protection would have much higher MPA costs (e.g. $4.5 billion for the developing world) but much lower opportunity costs, to the point of fisheries actually benefiting in the future. Since lower protection also compromises on biodiversity goals, our results show the trade-offs that political decisions need to consider beyond COP15. More generally, the unusually large opportunity costs show how marine contexts generate very different economic issues from terrestrial ones, by attempting to protect a common pool resource area that envisages no automatic market compensation for income lost to conservation.

---

Marine protected areas (MPAs) currently cover 8% of the ocean^1^, barely one quarter of what is needed to preserve marine biodiversity^2^. In response, Draft Target 3 of the Global Biodiversity Framework (GBF) has proposed expanding coverage to 30% by 2030^3^, abbreviated as “30×30”. However, there are serious concerns about the potential economic costs of such ambitious improvements in protection. First, governments can reasonably assume that a quadrupling MPA systems implies a large increase in the corresponding management budget, plus a one-off cost for creating so many new MPAs. Second, increased restrictions (e.g. fishing restrictions) could imply significant losses for marine livelihoods, economic output and food security.

The difficulty is that the size of these costs remains largely unknown, not least because the 30% proposal itself cannot yet include details about where or how it would be implemented in practice. Governments are therefore being asked to commit to a major policy change without knowing its cost - an issue we refer to as the “blank cheque” problem. Financial and economic costs are one of the major points of debate and disagreement at international environmental summits^4–6^, including the Conferences of Parties (COPs) of the Convention on Biological Diversity (CBD). In the absence of concrete information, this debate will revolve around imagined costs and losses, which can cause conflicts that are difficult to resolve and impede progress on policy. Three fundamental types of (unknown) cost can be defined. The first two are the direct costs of protected areas (PAs), namely the “management” cost (the annual cost of managing PAs) and the “establishment” cost (the cost of creating new PAs)^7–9^. The third type is the “opportunity” cost^10^, defined as the lost economic output resulting from greater use restrictions in PAs (although there can also be benefits e.g. from recovering fish stocks or enhanced tourism income in MPAs^11–13^). Most discussions of cost focus on the direct costs^8,14–16^, which are likely to be those paid by conservationists or environment ministries.

However, we argue that governments are more likely to achieve meaningful debate and implementation of ambitious CBD goals if they are given information about both types of cost together. Before all else, natural and social justice demands that environmental decision-makers consider the impacts of large policy changes on external parties, especially on the most vulnerable (including, for marine 30×30, the many small-scale fishers or SSFs who depend on exploiting the ocean to maintain marginal livelihoods^17^). More pragmatically, the final cost to governments may well include some additional budget to mitigate the negative impacts of greater protection on industries and communities, and so a focus on direct costs only is misleading. Even if a commitment is made to 30×30 with such limited information, implementation of the commitment may be weak or highly conflictive if it is later found that there are large opportunity costs. The draft GBF already contains language around permitting exploitative use of PAs that reflects an intuition of such opportunity costs^3^, but the vagueness of information about what those costs might be can translate into vagueness of protection, potentially helping neither biodiversity nor human communities in an efficient way.

Here, we estimate the full range of costs for the expansion of marine PAs to 30% coverage. We first develop three scenarios of 30×30 that reflect different balances between the priorities of conservation, the fisheries industry, and small scale fishers in particular. We develop new models that accurately predict the management costs of entire national MPA systems and apply them to estimate 30×30 management costs, also adding estimates of establishment costs. Simultaneously, we estimate the opportunity costs of the 30×30 target to the fisheries sector, under multiple sub-scenarios of future climate change and future approaches to making fishing effort more sustainable. Finally, we discuss the how marine conservation costs need to be approached in a fundamentally different way from the more commonly-studied terrestrial ones.

## Estimating costs of national MPA systems

Existing models of management and opportunity costs are unsatisfactory for resolving the blank cheque problem in several ways. For management costs, the most widely-used model is the one developed by Balmford et al.^15^ that predicts the cost of individual MPAs based purely on their size. But if one wishes to apply the Balmford model to a national MPA system, one first has to make a complex set of assumptions about how big each individual MPA will be, which is both time-consuming and uncertain^15^. More importantly, the assumption chosen can change the cost estimate by over 500% (an error that increases with the coverage desired); for example, Brander et al^18^. applied Balmford’s model to the same question (the cost of global 30% coverage) but derived a set of answers that were generally larger than those of Balmford et al. by hundreds of percent. The Balmford model also includes no estimate of central system costs. What is needed is a model that simply predicts national system costs directly, allowing rapid costing for proposals such as 30×30.

To create a costing algorithm for national MPA systems, we took empirical data from where governments or their agents have reported the “optimal” budget levels for their national MPA systems (defined as the funding needed for the system to achieve its goals fully), and then developed a mixed negative binomial regression model that predicts these budget needs (per hectare) (Methods). We used two different statistical estimation methods and these identified two possible best fitting models, based on information theoretic approaches (Methods). The first model (using the adaptive Gauss-Hermite quadrature approximation) contained terms for the national MPA system size, GDP per capita in the areas adjacent to the national system, and international tourist arrivals per capita (table 1, Methods). The second model (using the Laplace approximation) added, to this same set of terms, the ratio between the mean fisheries catch around the borders of the MPA system and the fisheries catch. Budget need per hectare went down as system size increased, reflecting well-known economies of scale^7,8,14,15^. Budget need also increased as GDP per capita in adjacent areas increased and (in the second model) as the relative fisheries catch increased, with the likely mechanism being that cost increase as pressure on the MPAs increases. Finally, budget need went down as the level of tourism increased. This was different from a result found in Australian MPAs^7^ where costs rose with visitor numbers. We hypothesise that the difference is due to the fact that our dataset contained many developing countries, where lower economic capacity and shortfalls in funding and staff can lead to some private tourism operators taking on many of the *de facto* management roles, reducing the perceived cost to state as the likelihood of tourism increases^19^.

**Table 1.**
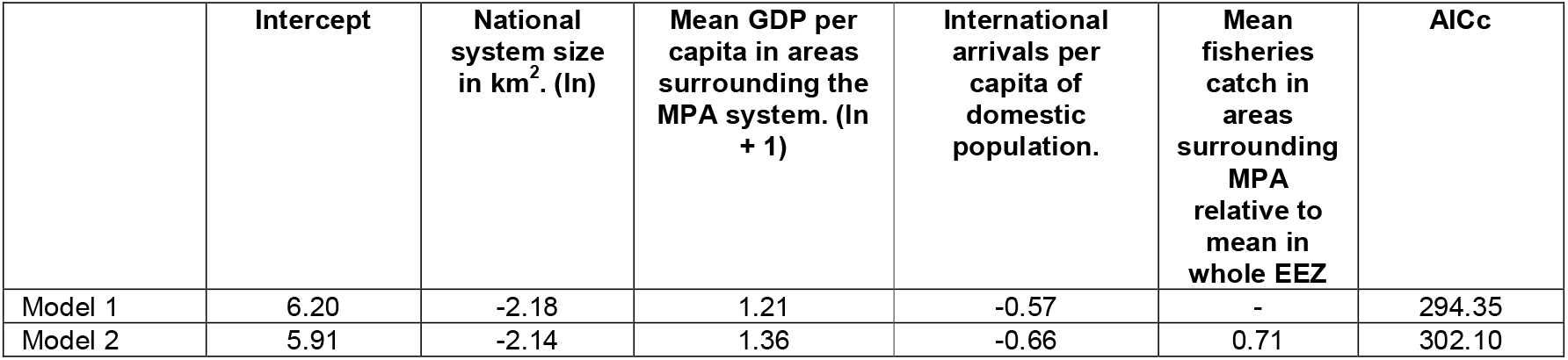
Statistical models predicting the optimal budget per hectare for a national MPA system. Budget need was input at purchasing power parity and ln-transformed. All predictor variables were z-standardized. Values indicate the model coefficients. See supplementary material for similar results using a different estimator software. The regression model was a mixed effects generalized linear model with negative binomial errors, a log link and a random intercept term for region.

|  | Intercept | National system size in km<sup>2</sup>. (ln) | Mean GDP per capita in areas surrounding the MPA system. (ln + 1) | International arrivals per capita of domestic population. | Mean fisheries catch in areas surrounding MPA relative to mean in whole EEZ | AICc |
| --- | --- | --- | --- | --- | --- | --- |
| Model 1 | 6.20 | -2.18 | 1.21 | -0.57 | - | 294.35 |
| Model 2 | 5.91 | -2.14 | 1.36 | -0.66 | 0.71 | 302.10 |

With a regression model in hand, the standard next step is to apply the model to predict costs of a larger PA system^8,15^. However, a system expanded to 30% coverage is likely to include a much larger proportion of offshore areas (e.g. beyond the 12 nautical mile limit) than the current MPA systems on which the model was parameterised. Offshore MPAs mostly require management of large industrial vessels that have remote tracking systems for monitoring and control^20^, presenting a very different cost context from inshore MPAs that can require intensive management of numerous small fishing vessels and other users. We therefore developed a further, novel costing model for offshore MPAs, based on observed costs of remote tracking and long-distance patrol vessels (Methods).

## Creating scenarios to allow costing of draft policy goals

MPA management costs depend on MPA location and governance arrangements but for policy proposals such as 30×30, neither of those has yet been decided at the moment of initial policy debate. To generate a rapid estimate of costs in the absence of such information, we created three 30×30 scenarios, rooted in biodiversity need but representing a range of political options about the strictness of protection (Methods, Supplementary Material). Since existing commitments to 30×30 largely interpret the target as 30% coverage for each country, we used that interpretation in our scenarios. Scenario 1 assumes that all MPAs in each country are no-take (i.e. no fishing); scenario 2 makes 50% of each country’s MPA area no-take and the other 50% mixed-use (i.e. some sustainable fishing is allowed); and scenario 3 uses the 50% mixed-use rule inshore (where small scale fishers operate) but assumes no-take rules in all MPAs offshore (where industrial fleets principally operate).

For each scenario, we divided each national system into an inshore component and an offshore one, varying the definition of these two areas to bracket a range of costs (Methods). To estimate management costs, we applied the predictive regression model to the inshore component and the offshore costing model to the offshore component. To estimate establishment costs, we took a recent model of cost per MPA creation^21^ and multiplied this cost by the likely number of new MPAs in each EEZ (Methods). We aggregate our estimates of all costs by World Bank income group because our intention is to give a broad sense of likely budget needs, rather than controversially suggesting that a budget is appropriate for any particular country before its government has even debated or chosen how to implement the GBF. The management cost estimates for high-income countries are less certain (Methods) so we report values for developing (non-high-income) countries separately, and then extend these to global values by adding the less certain estimate for high-income countries.

We found that the total annual finance needed to manage a 30×30 MPA system was $1.9-$4.5 billion for developing countries and $7.9-$14.4 billion globally (table 2). The average (median) cost per country ranged from $0.5-$0.7 million/yr for low income countries, to $2.0-$4.4 million/yr for lower-middle income countries, and up to $74.4-$120.1 million/yr in high income countries (figure 2, supplementary table 4). Our mixed-use approach (scenario 2) was considerably more expensive than a 100% no-take system (scenario 1). For example, scenario 2 was more expensive than scenario 1 by a factor of 2.4 in low income countries and 8.2 in lower-middle income countries. Establishment costs were much lower: the total additional budget needed would be $2.91 million for low-to-middle income countries (or $12.6 million if amortized at 5% over 30 years) and $5.7 million globally (or $24 million if amortized at 5% over 30 years) (table 3). This is equivalent to a mean annual payment over 30 years of $21,232 per country for low-to-middle income countries (or $91,764 at 5% interest) and $32,732 per country globally (or $141,465 at 5% interest). We caution that the global figures may be underestimated because some high income countries may have unusually large establishment costs^22^.

**Table 2.** Total estimated annual management cost to expand to 30% MPA coverage. Values show sum of costs for all countries in each World Bank income group, in millions of U.S. dollars (constant 2015 values) and using the mean of the 12 cost estimates for each country. scen = scenario,

| Income Group | total cost (\$ millions) | | |
| --- | --- | --- | --- |
|  | scen. 1 | scen. 2 | scen. 3 |
| Low | 13.6 | 32.1 | 22.0 |
| Lower-middle | 156.9 | 1285.9 | 257.6 |
| Upper-middle | 1758.1 | 3185.9 | 3161.9 |
| High | 6003.7 | 9855.8 | 8848.1 |
| All low- and middle-income | 1928.6 | 4503.9 | 3441.5 |
| . |  |  |  |
| Global | 7932.3 | 14359.7 | 12289.5 |

**Table 3.** Annualized establishment cost to expand to 30%, by World Bank income country group. Values are in millions of constant 2015 US dollars and show the median, 25th and 75th percentile for each income group; scen = scenario, min = minimum, max = maximum. Amortization calculations assume 5% annual interest on repayments over 30 years.

|  | median<br>cost | 25th<br>%ile | 75th<br>%ile |
| --- | --- | --- | --- |
| Income<br>Group | scen. 1 | scen. 1 | scen. 2 |
| Low | 0.004 | 0.003 | 0.077 |
| Lower-middle | 0.084 | 0.004 | 0.124 |
| Upper-middle | 0.110 | 0.026 | 0.164 |
| High | 0.010 | 0.006 | 0.242 |

## Projected impact on fisheries

To project possible opportunity costs, we input our three MPA scenarios as spatial areas of total or partial fishing restriction into two MEMs (BOATS and EcoOcean^23–25^), which estimate fish abundance from oceanic net primary production and sea surface temperature, then simulate future fishing effort, catch and value as a function of fish abundance and economic parameters. We defined the economic impact as the difference between each 30×30 scenario and a reference baseline of no further MPA expansion (in terms of both catch weight and catch financial value). The models calculated this impact yearly from 2025 (the first year of creation of new MPAs in the model) until 2100, under three alternative Marine Socioeconomic Pathways (MSPs): MSP1, MSP3 and MSP5. Each MSP made different assumptions about the severity of future climate change (the representative concentration pathway or RCP), the sustainability of future fisheries management practices and the joint impact of these on fish stocks and fishing patterns. MSP1 assumed RCP2.6 (i.e. very limited global warming) and a reduction of effort; MSP3 assumed RCP7.0 (higher global warming) and a steady increase in effort; and MSP5 assumed RCP8.5 (very high global warming) and unchanging effort (supplementary table 1 and 2). The two MEMs implemented “sustainable fishing” in mixed-use MPAs in different ways: BOATS reduced overall fishing intensity to sustainable levels for all fishers, whereas EcoOcean interpreted sustainability as permitting SSFs to fish freely, banning industrial fleets. For details, see Supplementary Material.

We report opportunity costs for MSP3 in the main text and for other MSPs in supplementary figures 1 & 2. For MSP3 in BOATS, catch values declined in all scenarios in the years immediately following 30×30 implementation, then started to recover. In scenarios 1 and 3, both of which have large proportions of no-take MPA, catch values remained below the reference baseline until the end of the century (figure 3). However, in scenario 2, which allows some sustainable fishing in MPAs, catch values after ~2040 rose to levels higher than the reference baseline. This economic benefit to fishers corroborates other studies that suggest that MPAs may promote fisheries stock recoveries (which could be exploited directly in mixed-use areas)^11,26^. Future work will incorporate estimates of the stock spillover that is observed to occur in a <200m strip along the border of fully no-take MPAs ^26^, a phenomenon not currently modelled by the MEMs. Opportunity costs may therefore be overestimated (although a 200m MPA-system border represents a very small fraction of total fishing area). Changing the assumptions about future fishing effort and climate change (in MSP1 and MSP5) did not affect this broad pattern but did alter the size of the projected changes (Supplementary Material).

Since EcoOcean effectively banned industrial fleets from all MPAs in all scenarios (varying the level of permitted access for SSFs only), the industry-wide impact for all scenarios was similar to the no-take scenario 1 for BOATS. We therefore used EcoOcean to study the specific impact on SSFs. This was highly variable, with some countries seeing SSF catch losses and others catch gains (Supplementary Material). The most likely mechanism explaining the gains is reduced competition with industrial fisheries, along with the same stock recoveries as before. Countries experiencing SSF gains included those where fishing was an important economic activity, such as many Pacific, Indian Ocean and Caribbean islands. However, there was not a clear economic difference between countries that gained and countries that lost catch.

## Implications for the financing of MPA expansion (30×30)

The managers of existing PAs already report that the funding they receive is insufficient to achieve their basic goals, with 50% shortfalls often reported^27,28^. Implementing the 30×30 target, without committing the finance to make it the MPAs effective, would result in a lose/lose situation. Conservation costs would go up, yet without meaningfully reducing biodiversity loss. Our study helps avoid this eventuality in two ways. First, it shows what the actual PA budget need is likely to be, so that governments can commit to the goal in full knowledge of the likely (approximate) cost. Second, it shows that the cost for MPA management is particularly modest: for example, the annual cost estimate for the entire developing world is $1.9-$4.6 billion per year (management and amortised (at 5%) establishment costs combined). Seen from this perspective (of direct costs only), a marine 30×30 seems highly feasible, especially if it receives some international support.

However, our study shows that in the marine context, the major cost can actually be the opportunity cost (corroborating Brander et al^18^.). If unaddressed, these large marine opportunity costs would risk 30×30 being unimplementable politically and practically^29^. MPA creation could be seen as particularly unjust because the government and international community is paying such a modest cost, and yet the hidden (opportunity) cost to local users - many of whom may be economically vulnerable - would be several times larger (a similar situation is faced by non-landowners who derive their livelihoods from terrestrial areas that become PAs). This negative impact will become even more important as the blue economy rows rapidly in size^30^.

Two contrasting approaches could address opportunity-cost issues. The first to allow exploitation of some parts of each national MPA system. Fishing currently occurs in 94% of MPAs^31^, a very high incidence that undermines marine biodiversity conservation^31^. Any example of a more balanced compromise than 94%, bringing reasonably positive outcomes for both fishers and conservation, may encourage decision-makers to implement greater strictness of protection in future. Although it was not our aim to define any “best compromise”, it is encouraging that our main compromise scenario (scenario 2) generated a small economic benefit for fishers, while giving full no-take protection to 50% of the MPA system and constraining fishing strongly in the other 50%. Detailed local planning of MPA zones could allow positive fisheries outcomes at even higher no-take percentages^29,32^.

The second approach is to accept a biological need for a fully protected (no-take) system and offer some form of financial mitigation to those suffering opportunity costs. In the extreme, a financial mitigation approach could be used with a 100% no-take MPA system that maximises biodiversity benefit (our scenario 1). Our results show that in that case, the annual mitigation spending needed may be several times larger than the management costs. For example, the median foregone catch value for a low or lower-middle income country is approximately $20-30 million per year for scenarios 1 and 3, whereas the corresponding management costs are only $0.5-$2 million (figure 3, supplementary 3) Economic losses do not necessarily need exact compensation but some form of mitigation, whether from subsidy, livelihood support and retraining, or other mechanism, is likely to be required for MPA expansion to be politically acceptable. The broader implication is that when costing marine PAs, it is particularly important to consider both direct costs and opportunity costs in a broader budget-need estimate.

More generally, our results emphasise that cost approaches honed on a largely terrestrial PA estate are misleading for a marine context. On land, PA costing models generally include the cost of purchasing new conservation land from tenure-holders^,29^. That purchase cost is generally high (approximately 50 times the annual management cost^33^) because purchase prices must compensate the tenure-holder for the future stream of income that s/he would expect otherwise^8,33^. In other words, the establishment cost incorporates much of the opportunity cost and allocates it to conservation authorities (such as the environment ministry). The oceans are a commons, and so ocean-users affected by MPA creation lack this tenure-based compensation. The lack of compensation for marine users has a particularly severe impact because new MPAs can easily be created on existing fishing grounds, making the loss of income immediate. New terrestrial PAs, on the other hand, would more often target unexploited land, so protection there does not imply an immediate, widespread loss of production.

If MPA implementation is to be just and politically feasible, marine conservation managers may need to bestow on the commons a set of economic rights similar to those enjoyed by tenure-holders of targeted conservation land. Including some form of financial support in the estimate of marine budget needs also has an element of social-justice logic: with MPAs, governments are essentially saved from paying very large establishment costs, precisely because there is no marine framework for compensating tenure-holders for lost income. We do not mean to suggest that governments should add up support and management costs and simply choose the cheapest option. Nor do we recommend any of our scenarios in particular, noting simply that they represent different approaches to trade offs that sovereign governments and their rights-holders and stakeholders must decide between. However, our models can give a broad sense, in advance of any decisions made, of what the likely economic consequences are. Some mixture of financial mitigation approaches and permitted-fishing approaches in MPAs may eventually be needed, depending on local conditions and biodiversity needs.

In conclusion, the management costs, opportunity costs and biodiversity benefits of MPAs all trade off against each other. A highly protected system of 30% MPAs would require very modest MPA managements budgets, but any savings could be dwarfed by the size of the financial support needed. Allowing some fishing in MPAs can greatly reduce opportunity costs (and therefore financial support needs) and even boost fisheries incomes, but at the cost of higher management budgets and reduced biodiversity protection. Mitigating the opportunity costs, while making protection stricter, is an alternative that respects the particular economic reality of the ocean economy: that fishers, as users of a commons, can suffer immediate income losses after MPA creation, yet without enjoying the right to compensation that owners of land enjoy. At a time when rights-based approaches to conservation have come to emphasise the importance and justice of PA impacts on non-landowners^34^, a logical extension is that users of the marine commons should be granted the same consideration.

## Supplementary Material

### Methods

To generate a costing algorithm for national MPA systems, we first collected data on the empirical budget needs of existing systems (n=30, indicating the difficulty and rarity of estimating optimal budgets), where “budget need” refers to the budget that should be allocated if the system is to achieve its objectives, not the current (often inadequate) budget. Following several previous authors^8,14,15^, we chose to analyse the budget need per hectare. Data on budget needs came from a variety of literature sources but most represent the estimate of a high-level, usually governmental agency of the finance needs of an entire system or in a few instances, a major subsystem within the national system. Many of the sources are confidential, due to the sensitivity of government financial data, and so are available upon considered request and subject to a confidentiality agreement. Budget need is often assessed at two levels of adequacy: “basic” (the minimum budget needed to maintain core management activities adequately) and “optimal” (the estimated cost of managers fully achieving the protected areas’ goals)^14,35^. Sources reporting budget needs reported optimal rather than basic needs more often, and so we analysed optimal needs.

To create a costing algorithm that could predict finance needs in both no-data countries and for expanded MPA systems, we first aimed to identify a statistical model that could predict the empirical data with a high level of accuracy. We compiled a set of variables that are theoretically expected to influence finance needs for MPA systems. These variables are listed in supplementary table 1.

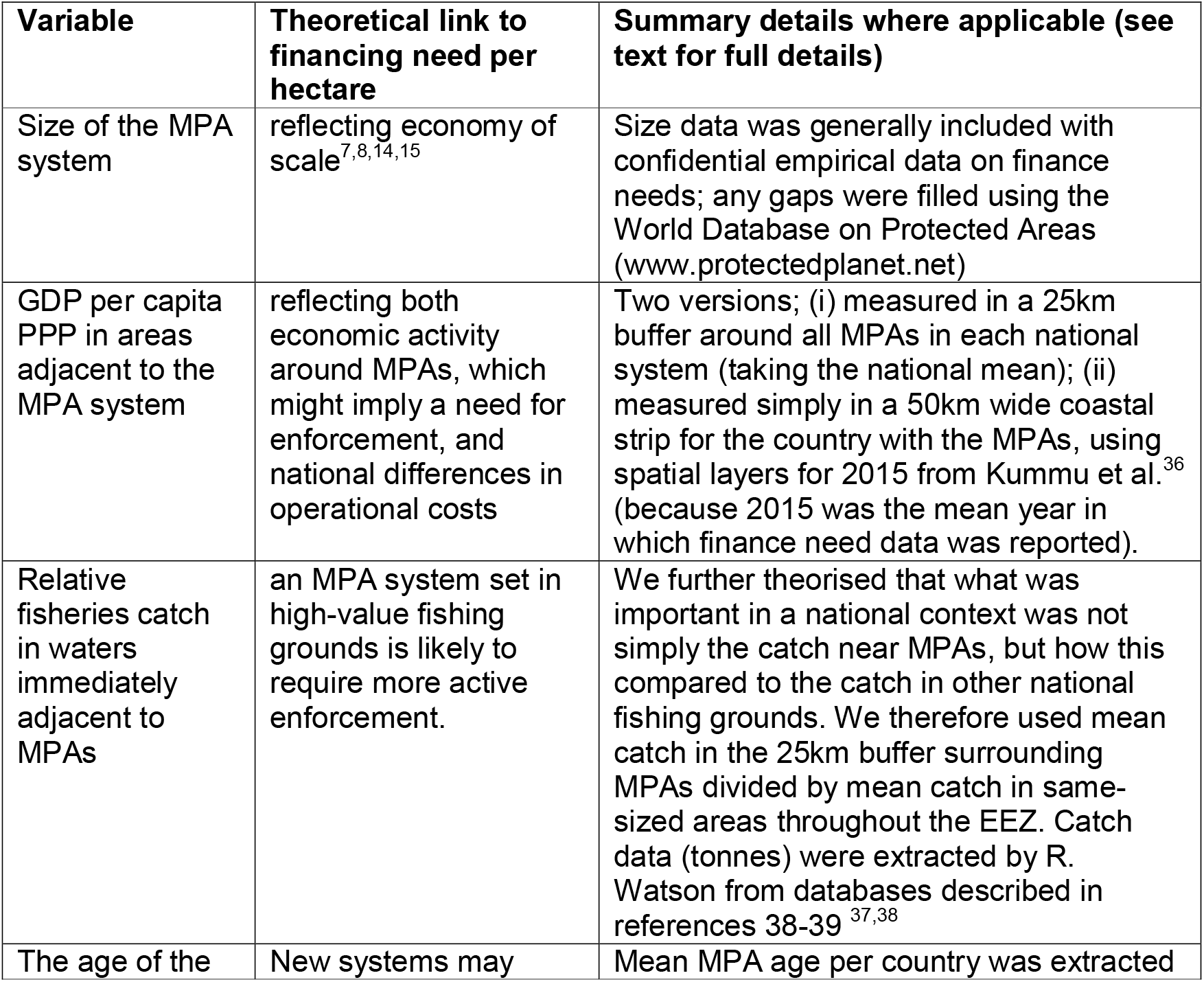
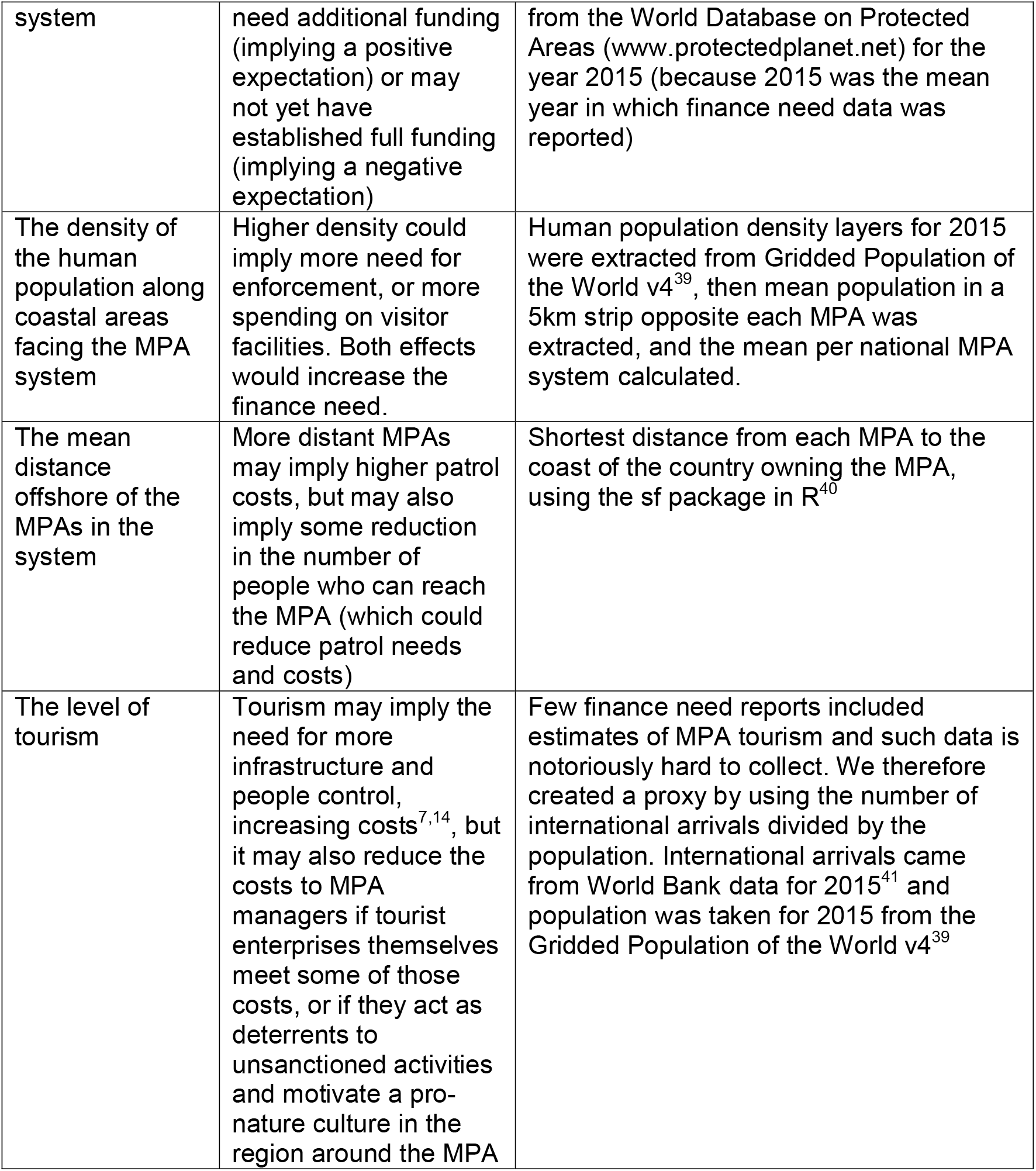

GDP and population size in coastal areas near MPAs is often calculated by buffering out MPAs to a fixed distance and then analysing GDP or population inside the buffer. However, using a fixed buffer size can create a situation where a nearshore MPA has a buffer that stretches far inland, whereas a further-offshore MPA has a buffer that only just touches a few metres of coast (and can also cause distance offshore and the buffer values to interact non-randomly). We preferred to have the buffer reach the same distance inland in all cases and then create an independent distance variable, and so we created customised buffers for the MPA system by adding 5km to the offshore distance (essentially creating a 5km strip of coastal land opposite each MPA). We then extracted data on GDP/capita PPP^36^ and human population size and density^39^ for 2015. However, we did follow the logic of previous authors by capping the buffer size at a maximum of 50km, firstly because we were using a different costing algorithm for MPAs that are far offshore (see Offshore Costs), and secondly to reflect an intuition that the impact of coastal populations would dwindle to a low level in MPAs that are far offshore.

For St Kitts and Nevis, calculating the GDP in a set of coastal strips opposite the MPA system produced NAs in spatial analysis. The calculation itself is also an onerous step if the costing method is to be used to cost future MPA expansion proposals. We therefore tested whether GDP per capita for each country’s 50km coastal strip was a reasonable proxy of GDP per capita in the customised buffers. We found that the simpler measure (the 50km coastal strip) was an accurate predictor of the more complex one (R-squared = 0.912) but also tended to overestimate the buffer-based result, for the clear reason that a buffer contains a seaward side in which there is no nearby coast and GDP is therefore zero, reducing the mean value in a way that does not occur if one only measures the coastal strip values. We therefore adjusted coastal GDP by applying the regression relationship between itself and the empirical data on buffer GDP (which has the effect of reducing the coastal strip-based GDP value).

For all variables except system size (which was extracted at national scale), we calculated national values as the mean of the values for all individual MPAs in a national system. All predictor variables were z-standardised prior to regression analysis and natural log transformations were applied to the system extent and to GDP.

Due to the hollow-curve, non-negative frequency distribution of our finance-need data, our regression analysis used mixed generalized linear models with a negative binomial error structure, a log link and a random effect for region (via the intercept), using two different fitting packages in the R software environment: lme4 (with the glmer.nb command)^42^ and glmmadapative^43^. We used an information theoretic approach to select best-fitting model or models, ranking by delta AICc (Akaike Information Criterion corrected for sample size)^44^. Since information theoretic approaches select the best model from those provided by the researcher but are mute as to the explanatory power of the terms included, we also extracted p values from the best fitting regression models and checked whether p<0.05. Finally, as a measure of goodness of fit for a mixed negative binomial model (where R-squared values are not naturally estimable), we calculated and plotted the correlation between the predicted and observed values (figure 1). Diagnostic plots for regression analysis were also examined and these suggested no problems with the best-fitting models selected.

**Figure 1.**
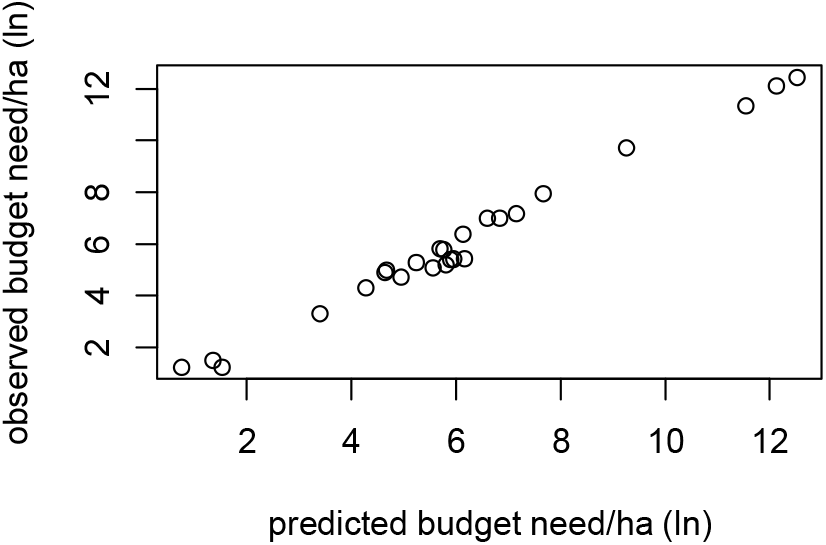
Comparing observed data on MPA budget need per hectare against the predictions of the statistical model.

**Figure 2.**
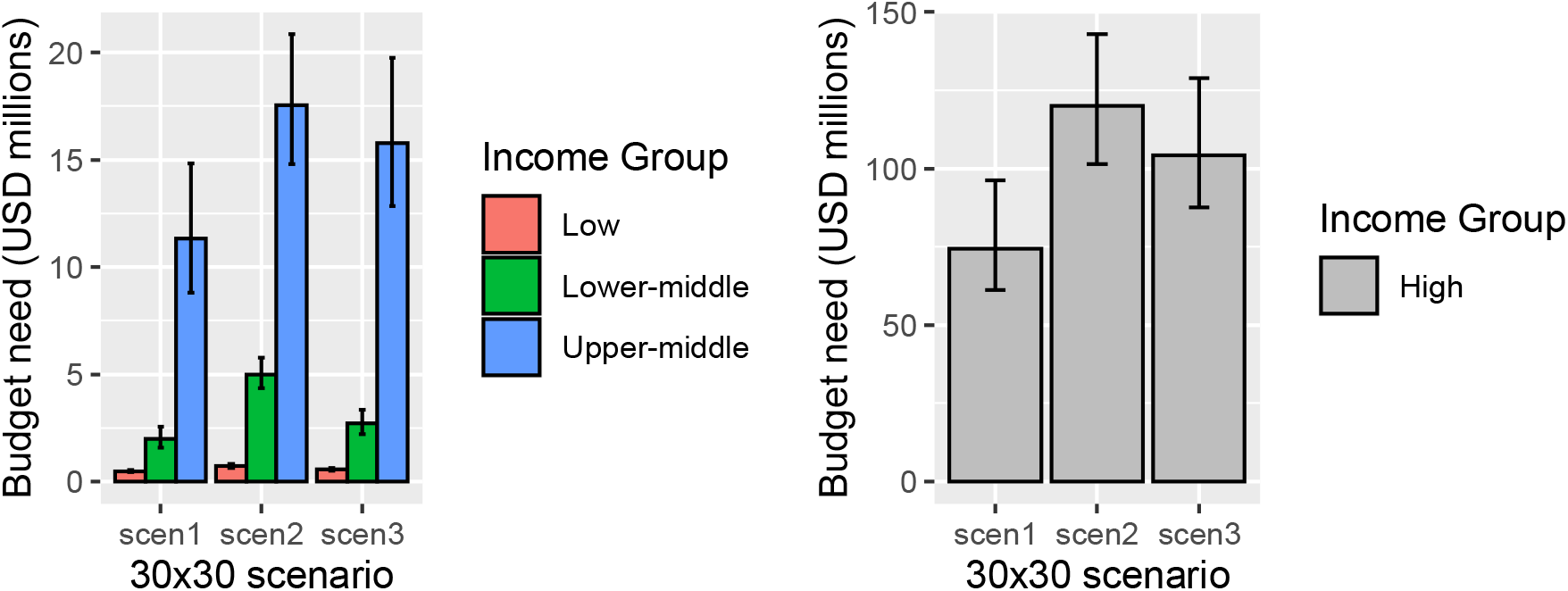
Median annual management costs of MPA system with 30% coverage, under different scenarios. Results show the median cost per country in each income group; error bars show the minimum and maximum for any country in that group (see Methods). Values are in constant 2015 US dollars.

**Figure 3.**
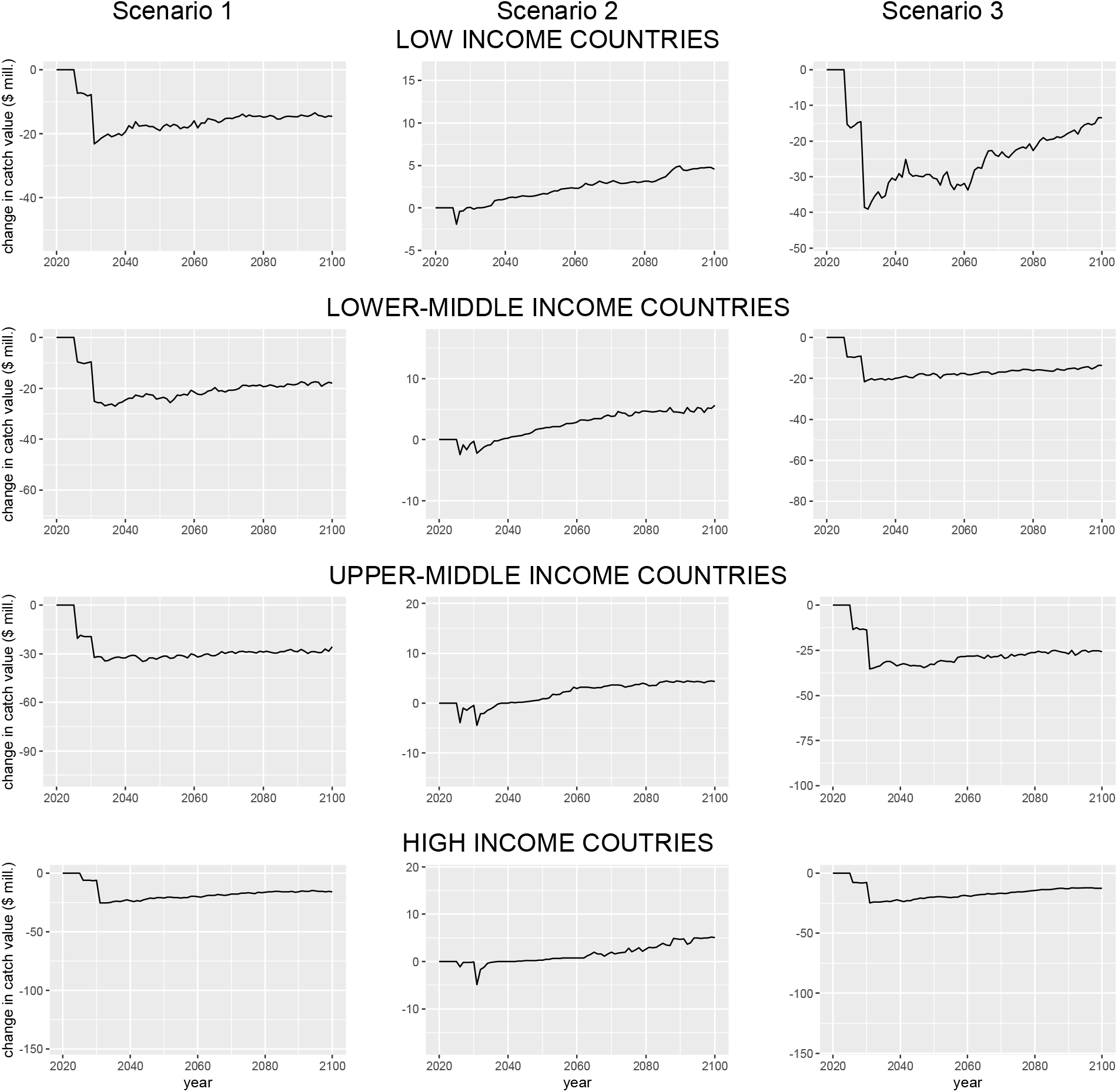
Projected change in the value of fisheries catches (relative to the reference baseline) under the mid-range marine socioeconomic pathway (MSP370). Line shows the median and shaded area shows the interquartile range of projected opportunity costs for countries in each income group. Note the different scales for the different scenarios. Values are in constant 2015 US dollars.

Empirically, MPA systems at the time of the budget-need reports tended to be predominantly placed in inshore waters and rarely occurred in far-offshore parts of the EEZ (e.g. in our data, a mean of 47% of the total MPA area lay inside the 12 nautical mile limit and thus inside only 11% of the EEZ). By implication, any costing model parameterised upon that data is best suited to predicting costs for more inshore areas. However, expansion of MPA coverage to 30% of the EEZ is likely to create several new protected zones that are considerably further offshore (for example, our 30×30 scenarios (below) suggested a median of 77% of the MPA area would be outside the 12 nautical mile limit). This creates a problem of applicability for the regression model because management of an offshore area differs notably from management of an inshore area, implying that offshore areas could also have very different costs. Specifically, offshore areas are likely to be mostly under threat from large industrial actors such as industrial fishing boats. Industrial fleets that can be in distant parts of the EEZ are most efficiently controlled by a combination of long-distance patrol vessels and remote monitoring technologies^20,45^, which combine tracking of ship courses, independently checked onboard CCTV cameras and sensors that detect fishing gear use (see also Waldron in preparation). Nearer the shore, actors such as small-scale fishers and tourists require more intensive management and their vessels often lack the onboard technology that would allow remote monitoring.

We therefore decided to split the costing estimate into two parts: an inshore component and an offshore component. The inshore component was estimated using the regression model. For the offshore component, we reviewed the management cost data from very large MPAs and supplemented it with data from some large offshore fishing grounds that required enforcement (Waldron in preparation). We extracted the size of the industrial fleets operating in each EEZ from Tidd et al.^46^ and calculated the combined cost of installing and administering remote sensing, remote monitoring via onboard cameras for that fishing fleet (with recordings independently verified by a paid professional), plus a patrol fleet similar to that used in fisheries and very large MPA enforcement to date (a range of three possible values assuming either one boat per 500,000km^2^, per 400,000km^2^ and 600,000km^2^.

#### Model Results and Application

One of the principal difficulties of costing a proposed target (such as 30×30) is that the costing must be done in advance of any specific information. Intuitively, the costs are likely to depend strongly on the spatial distribution and protection regimes of any future MPAs, and yet at the moment of policy creation, none of these details have been decided. One solution to this paradox is to take a set of scenarios of how the policy might reasonably be implemented and analyze each one. In this way, a range of costs can be estimated, which are assumed to be likely to bracket the eventual cost. Any estimated cost using this method cannot therefore be fully accurate, nor is it intended to be. Instead, the goal is to provide a range of likely costs and economic impacts, and to illustrate likely trade offs.

With this goal in mind, we created three scenarios for 30×30, plus a reference scenario of no further expansion of MPAs (i.e. MPAs remain static at their 2021 extent and strictness of protection). All scenarios begin from the current (2021) extent of MPAs (downloaded from the World Database of Protected Areas, www.protectedplanet.net), and then expand the system in each country to achieve 30% coverage of the EEZ. We divided the EEZ up into 1km squared cells and identified all cells already protected. Expanding from that baseline requires a ranking system and for this cell ranking, we used the raster map of marine biodiversity conservation priorities from Sala et al.^11^. The Sala et al. ranking system uses equal-area cells of approximately 3000km^2^, so we downscaled the original rasters to the 1km resolution and clipped to the borders of the EEZ. (Spatial EEZ outlines were taken from the World EEZ v11 product at marineregions.org). We then added 1km cells to the existing MPA system in order of their cell values. If, in the last iteration of this addition process, the cropped 3000km2 cell with the next highest value produced more 1km cells than were needed to achieve 30% coverage, we chose to preserve contiguity (since such a practicality was likely to reflect political and operational realities), by adding the 1km cells contiguously in a west-east direction until the 30% target was reached.

The basic spatial layer of cells extracted so far had no definition of the strictness of protection or access rules implied. Since part of our interest lay in the direct costs and opportunity costs associated with different compromises between biodiversity and fisheries livelihoods, we created three scenarios on the basis of the basic layer, differentiating between scenarios by varying the fishing effort permitted in different parts of the EEZ. Specifically, the scenarios were created to explore the differences in all costs between a biodiversity focused system that was 100% no-take; a system that allowed compromise between biodiversity protection and all fishers, including both industrial and near-shore/small scale ones; and systems that compromised in ways that would protect small-scale and near-shore fishers’ livelihoods but excluded large industrial fishing vessels from protected areas (to the nearest spatial approximation. For simplicity, we defined the fishing access permitted in the different scenarios by defining two protection levels, using a simplified structure similar to current categories of protection^47^. “High protection” areas were modelled to exclude fishing exploitation (similar to Horta e Costa et al.’s Fully Protected Area category^47^), and “medium protection” areas were modelled to allow a low/sustainable level of fishing carried out (similar to Moderately Protected in the same source). In BOATS, sustainable fishing in MPAs is defined as follows, irrespective of whether the fisher is industrial or small-scale: If fish populations are overfished in 2014, sustainable fishing is the effective effort level in 1974. If there was no fishing in 1974, sustainable fishing is half of the effective effort in 2014. If fish populations are not overfished by 2014, sustainable fishing is the effective effort in 2014. If there was no fishing in 2014, fishing is left unregulated. In EcoOcean, sustainable fishing definitions focused instead on often-vulnerable small-scale fishers (SSFs) and simply defined medium protection as allowing fishing by SSFs but not by industrial vessels. For management cost model, medium protection was defined following Ban et al^7^. where it is the simple difference between no-take and “not no-take” that drives management cost differences.

We then divided the selected MPA cells in each scenario into an inshore component and an offshore component for each EEZ. To avoid arbitrariness in the choice of where these two components occur in marine space, we created four separate definitions of the limit line between inshore and offshore. The first inshore/offshore limit line was based on the mean of the empirical MPA area contained inside the 12 nautical mile limit (47%, which we converted to a limit line at 2.12 times the 12 nautical mile limit). We also modelled a limit line sitting exactly on the 12nm limit (generating the smallest inshore extent); one lying 1.55 times 12nm offshore (based on the 66.67th percentile of the empirical distribution of the percentage MPA area inside the 12nm), creating the second smallest inshore area; and one lying 50 nautical miles offshore (a point at which small scale fishers are likely to be reduced to very low frequency^48^), where this last definition created the largest inshore area. These different limit-line definitions essentially represent a management decision about how far offshore it is worth continuing to apply (more expensive) inshore approaches. For example, the further offshore one goes, the fewer small actors are likely to be present (due to differences in feasible sailing distances and engine power). There will therefore be a point (a distance) offshore at which the managing agency decides it is no longer worth using intensive inshore management techniques and switches to more at-a-distance offshore techniques. We acknowledge that it may sometimes be possible to apply both intensive and at-a-distance techniques. However, simultaneous application of both approaches is unlikely to change the cost estimates materially because small inshore vessels generally do not carry the technology that allows them to be tracked and any larger vessels that do move close to shore will already have had their tracking costs factored in as part of the offshore model.

To translate different scenarios into spatial rasters for analysis, we divided the base 30×30 layer of selected cells up so that in each scenario, a different pattern of protection levels applied across the national system as a whole, without needing to make any assumptions about protection mosaics in each individual MPA but respecting the broader difference between inshore and offshore areas. Our range of approaches for protection levels was designed to explore trade-offs between the demands of national fishers, coastal livelihoods and conservation priorities (supplementary table 2). Thus, scenario 1 assumes no-take for the entire national MPA system, including those already existing; scenario 2 splits the national system 50:50 between medium and high protection; and scenario 3 has a more complex arrangement in which the compromise of medium protection applies only in the inshore areas where SSFs are likely to operate (supplementary table 2). For the Reference Scenario (MPAs already in existence), we defined high protection as all current MPAs classed as Fully/Highly Protected in Sala et al.^11^ (op. cit.), and medium protection as all other current MPAs. (We acknowledge that some current MPAs have insufficient protection to be classed as “medium protection” (for example, they may still be unsustainably fished), and this is often associated with very large funding shortfalls for their management^27^. The reference scenario itself should therefore be regarded as an ambition for the future, in which all current MPAs are fully funded and can achieve at least a sustainable level of exploitation (medium protection) and where desired, high protection.)

**Supplementary Table 2.** Scenarios used to create a range of illustrative costs for 30% MPA protection.

| Scenario | Description |
| --- | --- |
| Reference Scenario (extent of current MPAs) | Assumes no future MPA growth beyond what is currently in place. However, existing MPAs are modelled to receive the fully adequate level of funding. Existing MPAs that are not Fully Highly Protected are all modelled to achieve medium protection (sustainable exploitation). Existing Fully Highly Protected MPAs retain high protection status. |
| Scenario 1 (30% of EEZ) | Very stringent conservation: all 1km cells are given high protection, including upgrading of existing MPAs that are not Fully Highly Protected |
| Scenario 2 (30% of EEZ) | Full trade-off between fisheries and conservation. Both inshore and offshore, 50% of the expanded MPA area is modelled as high protection and 50% as medium protection (preserving classification of existing MPAs). NB in one of our Ocean Ecosystem Models (EcoOcean), high protection still allows artisanal fishing. |
| Scenario 3 (30% of EEZ) | Partial trade-off between fisheries and conservation. Inshore, 50% of the expanded MPA area is modelled as high protection and 50% as medium protection (with existing MPAs preserving their current classification). Offshore, all 1km MPA cells are modelled to have high protection, excepting that any existing offshore MPAs that are not Fully Highly Protected retain their existing classification. |

To apply the predictive regression model and extract inshore costs, we created a new dataset of system sizes for the 30% scenarios, breaking down the size into the inshore extent and the offshore extent and using the inshore extent only as the input to the model for prediction. This was done four times to capture the four alternative spatial definitions of inshore/offshore area that we had created earlier. We also calculated the area of each inshore and offshore area, in each EEZ, that would receive high and medium protection respectively. For the GDP per capita term, we input GDP per capita PPP in the 50km coastal strip of each country. For relative catch, it was difficult to robustly project the likely future catch in the areas around new MPAs post-2030 and how catches in other parts of the EEZ (i.e. not associated with MPAs) would change, so we simply applied a range of possible multipliers of the current relative catch and used them to sensitivity-test the outputs of the costing model (supplementary results). We did not attempt to forecast changes in the future ratio of international arrivals to domestic population, taking the simplifying assumption that it would remain constant.

To account for offshore management costs, we calculated the cost of one long-range patrol boat per 500,000km^2^ (varied to 400,000 and 600,000 to give a range of values) and the REM need (including satellite and radar costs, administrative costs and the costs of monitoring onboard video recordings) for all industrial fishing vessels in the EEZ, assuming that when MPAs occupy 30% of a country’s ocean, often in remote parts of the EEZ, then the entire fleet should be tracked and monitored. We therefore assumed that offshore areas would be fished by large vessels and took the number of large vessels in each EEZ from Tidd et al^46^. Future vessel numbers may vary but given the small impact this variation would have on the overall cost, we again took a simplifying assumption that they remained at the same level observed in the empirical data.

We further adjusted the predicted management cost of the inshore component to capture differences in cost between high and medium protection. To allow a range of uncertainty around this cost differential, we used three different estimates based on Ban et al.’s analyses^7^, in which experts and statistical models both suggested cost savings if operating a system with 100% no-take rather than a mixed system with 30% no-take. Experts suggested 20-year management costs would be 1.69 times higher for the mixed system and models suggested 1.96 times (with other values also possible). We therefore assumed, as our three alternative adjustments, that high-protection MPA systems would need annual management budgets that were, respectively, 51% (=1/1.96), 54.7% (=1/(1.96+1.69/2)) and 59.2% (=1.1.69) of the budget needs for medium protection systems. (We note that Balmford et al.’s MPA cost model^15^ did not include a significant effect for no take/extractive use allowed, whereas Ban et al. reports that both experts and a statistical analysis concurred that no-take MPAs have lower management costs). For the offshore component, we assumed that due to the remoteness and large distances involved, similar vessel tracking and patrol boat capacity would be needed across the range of access rules.

#### Establishment costs

Establishment costs are those costs associated with the initial creation of a new MPA. Two studies have extracted known costs and sought models to explain them: McCrea-Strub et al.^22^ found data from 13 MPAs across 6 countries and found the costs to be related to MPA size and the duration of the creation phase, with the size relationship strongly driven by the difference between a cluster of very small MPAs (largely in the Philippines) and three very large MPAs, two of which were in the USA. Binet et al.^21^ found data from 23 MPAs across 10 countries and found the cost to be independent of size. When attempting to model the creation costs for an entire national system which has not yet been designed, the Mcrea-Strub model obliges the researcher to assume the size of each new MPA in the unknown system, presenting the same problem as occurs when trying to apply the Balmford model for individual MPA management costs to predict the cost of an entire unknown system. Since the Binet model is size-independent and based on a larger sample, we used that for the present study. The Binet model does require an estimate of the number of MPAs in the system. To derive this number, we first assumed that the inshore part of the expanded system would be divided into multiple MPAs but the offshore part would be patrolled by a single long-range vessel assisted by remote monitoring, making it essentially a single unit. We then extracted the number and extent of inshore MPAs in all current national systems (using the 12nm definition of “inshore”), calculated the statistical relationship between number and extent of MPAs in existing data for 2019 (www.protectedplanet.net) and used that statistical model to estimate the number of MPAs expected in the inshore component of the 30×30 scenarios, rounding to the nearest integer then adding one for the assumed offshore unit. This calculation generated a mean of 15 new MPAs and a maximum of 81 new MPAs per EEZ.

#### Opportunity costs

The principal opportunity cost studied in MPAs is typically the possible loss of fishing revenues (foregone catch value)^18^ and we focus on that cost here. (We do not look at foregone mining and extractive activities). We projected fisheries opportunity cost by interpreting MPAs as a set of restrictions on fishing activity (again varying the amount of MPA that had medium protection and allowed some sustainable fishing) and then using MEMs (Marine Ecosystem Models) to project the likely changes in catch value that result. We used two MEMs (BOATS^24,25^ and EcoOcean^23,49^), since each model represent the many complexities of marine ecosystem functioning in different ways. BOATS was used to extract aggregate economic projections but does not disaggregate by Small Scale Fisheries (SSF) and Industrial Fisheries. We quantified the opportunity cost from BOATS as the difference between the projected value of the catch under each scenario (in each year) and the projected value of the catch under the reference scenario. EcoOcean does estimate SSF outcomes separately but given the uncertainties involved in estimating the precise dollar values of catch across a large number of small craft with limited record-keeping and the fact that some SSF catch effort may be intended for personal consumption rather than market sale, we quantified the SSF opportunity cost as projected percentage changes in catch between the 30×30 scenarios and the reference scenario.

The MEM outcomes also depend on assumptions about the level of ambition (and success) in future sustainability goals affecting the ocean economy more broadly, particularly with respect to fisheries management and climate change (RCPs or representative concentration pathways). To reflect this range of possibilities, three sub-versions of each opportunity cost projection were generated, each based on a different marine socioeconomic pathway (MSP), which describes a combination of a shared socio-economic pathway (SSP) to model trajectories in the sustainability of fisheries management (interpreted for the ocean system by Maury et al.^50^), plus a representative concentration pathway (RCP) to show an associated level of sustainability in climate change mitigation:

1. <u>MSP1 = SSP1+ RCP2.6</u> Fishing effort is steered back to sustainable levels thanks to strong management and long-term considerations. The effort in 1974 is defined as the sustainable baseline, and in projections effort should return to 1974’s levels by 2050. In EcoOcean: nominal effort changes. In BOATS: effective effort changes.
2. <u>MSP3 = SSP3 + RCP7.0</u> Fishing effort increases due to poor management and short-term priorities, but rate of technological progress is low. Rate of nominal effort increase is 1% yr-1, based on Rousseau et al. In both models, nominal effort changes.
3. <u>MSP5 = SSP5+ RCP8.5</u> Fishing effort is diversely affected by decreases in demand, poor management and high technological progress. This complex development is implemented by fixing effort at the 2015 levels. In EcoOcean, nominal effort is fixed. In BOATS, effective effort is fixed.

The complete set of model runs, accounting for both scenario differences and SSP/climate forcing options is shown in supplementary table 3.

**Supplementary Table 3:**
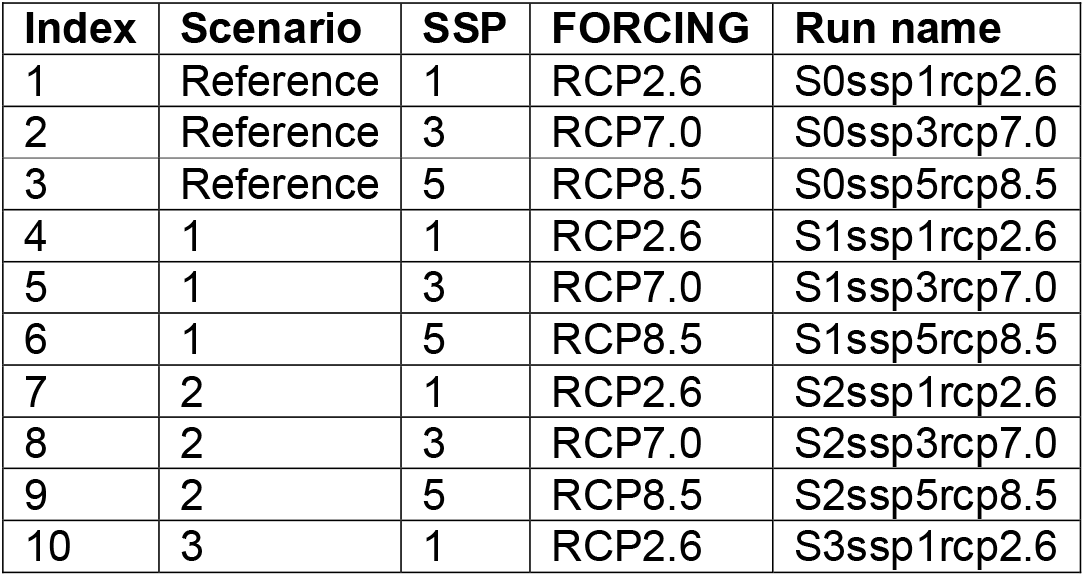
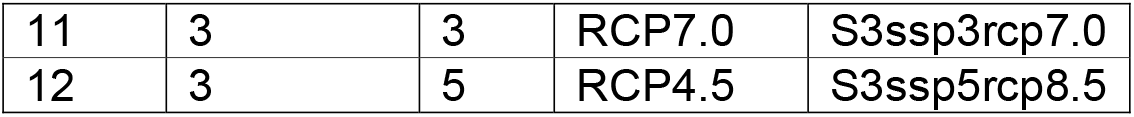
List of scenario runs for the Ocean Ecosystem Models.

| Index | Scenario | SSP | FORCING | Run name |
| --- | --- | --- | --- | --- |
| 1 | Reference | 1 | RCP2.6 | S0ssp1rcp2.6 |
| 2 | Reference | 3 | RCP7.0 | S0ssp3rcp7.0 |
| 3 | Reference | 5 | RCP8.5 | S0ssp5rcp8.5 |
| 4 | 1 | 1 | RCP2.6 | S1ssp1rcp2.6 |
| 5 | 1 | 3 | RCP7.0 | S1ssp3rcp7.0 |
| 6 | 1 | 5 | RCP8.5 | S1ssp5rcp8.5 |
| 7 | 2 | 1 | RCP2.6 | S2ssp1rcp2.6 |
| 8 | 2 | 3 | RCP7.0 | S2ssp3rcp7.0 |
| 9 | 2 | 5 | RCP8.5 | S2ssp5rcp8.5 |
| 10 | 3 | 1 | RCP2.6 | S3ssp1rcp2.6 |
| 11 | 3 | 3 | RCP7.0 | S3ssp3rcp7.0 |
| 12 | 3 | 5 | RCP4.5 | S3ssp5rcp8.5 |

MEMs were run at a resolution of one degree on a latitude-longitude grid of the oceans. Scenario designs were therefore upscaled from 1km to one degree, with parallel grid inputs showing the proportion of high protection and the proportion of medium protection in each one-degree cell. We note that this has the advantage of not imposing exact locations on the MPAs, in keeping with the need for sovereign countries in consultation with key stakeholders and rightsholders to retain control and flexibility in the final location of any MPAs. To reflect the time taken to create expanded MPA systems and the way that implementation of that goal is likely to be staggered in time, the expansion was implemented in the MEMs in two parts: half of the new MPA area was modelled as coming into existence in 2025 (by random draw), and the remainder in 2030 by linear interpolation.

As output, the MEMs projected the following economic outcomes at five-year intervals from 2020 to 2100: catch (mass), catch (value), catch per unit effort, and net catch value (profit). Five-year values were calculated as the average of the preceding five years. Overall, the two MEM model outputs (for BOATS and EcoOcean) were therefore harmonized to the highest degree possible.

#### Supplementary Results

**Supplementary Figure 1.**
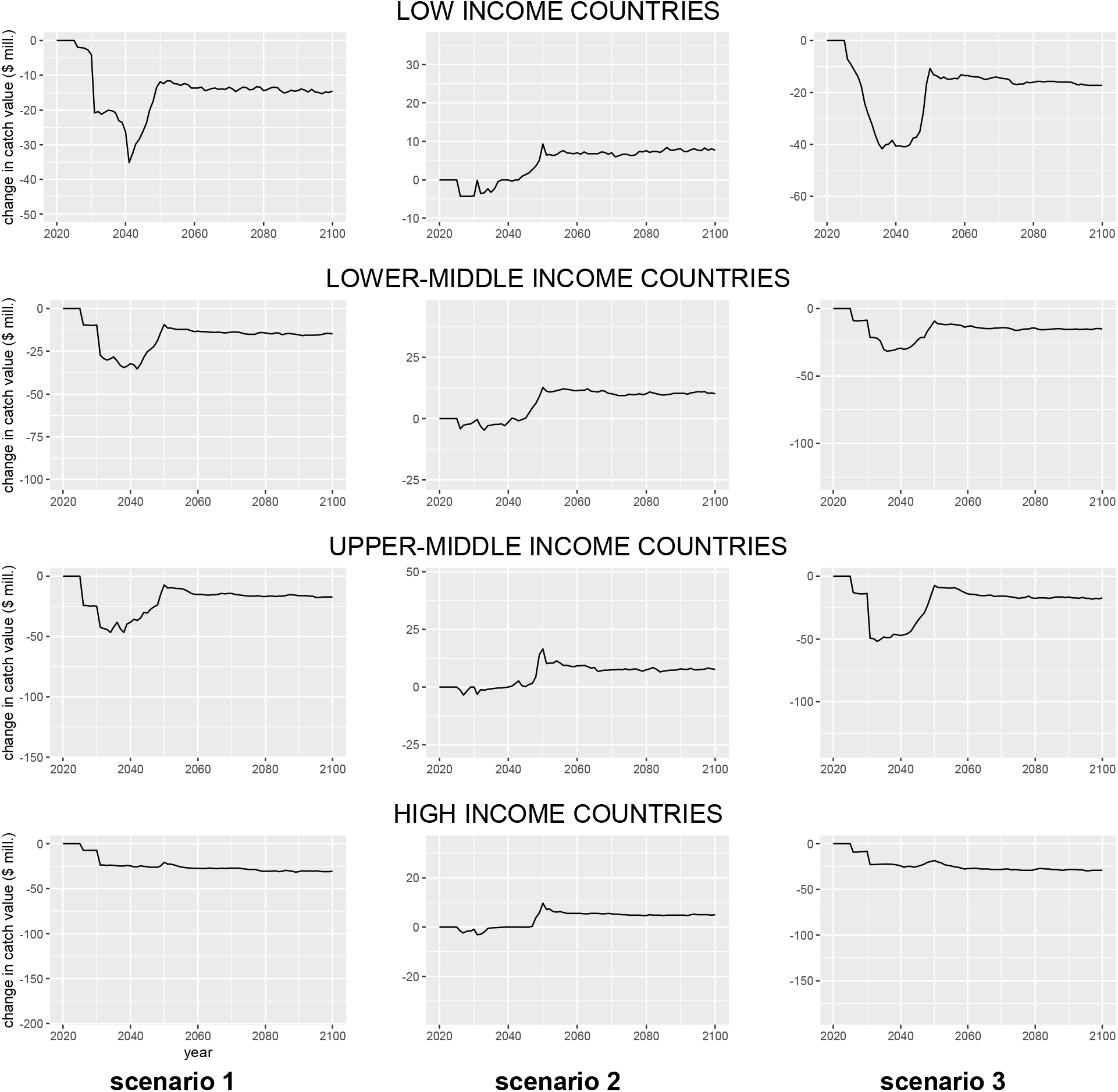
Projected change in the value of fisheries catches (relative to the reference baseline) for MSP 126. Line shows the median and shaded area shows the interquartile range for each income group and each scenario. Note the different scales for the different scenarios. Values are in constant 2015 US dollars.

**Supplementary Figure 2.**
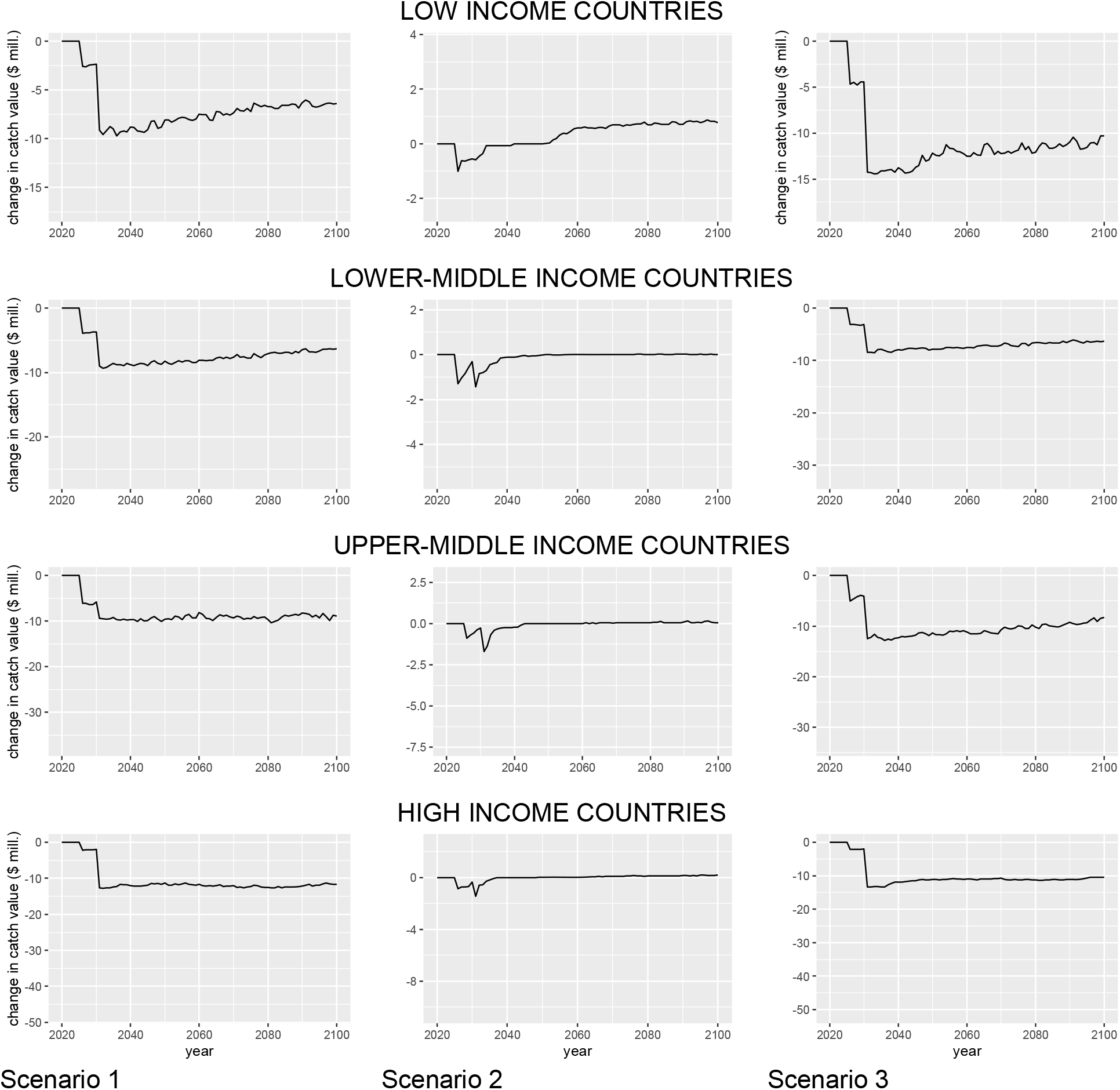
Projected change in the value of fisheries catches (relative to the reference baseline) under MSP585. Line shows the median and shaded area shows the interquartile range for each income group and each scenario. Note the different scales for the different scenarios. Values are in constant 2015 US dollars.

**Supplementary Table 4.**
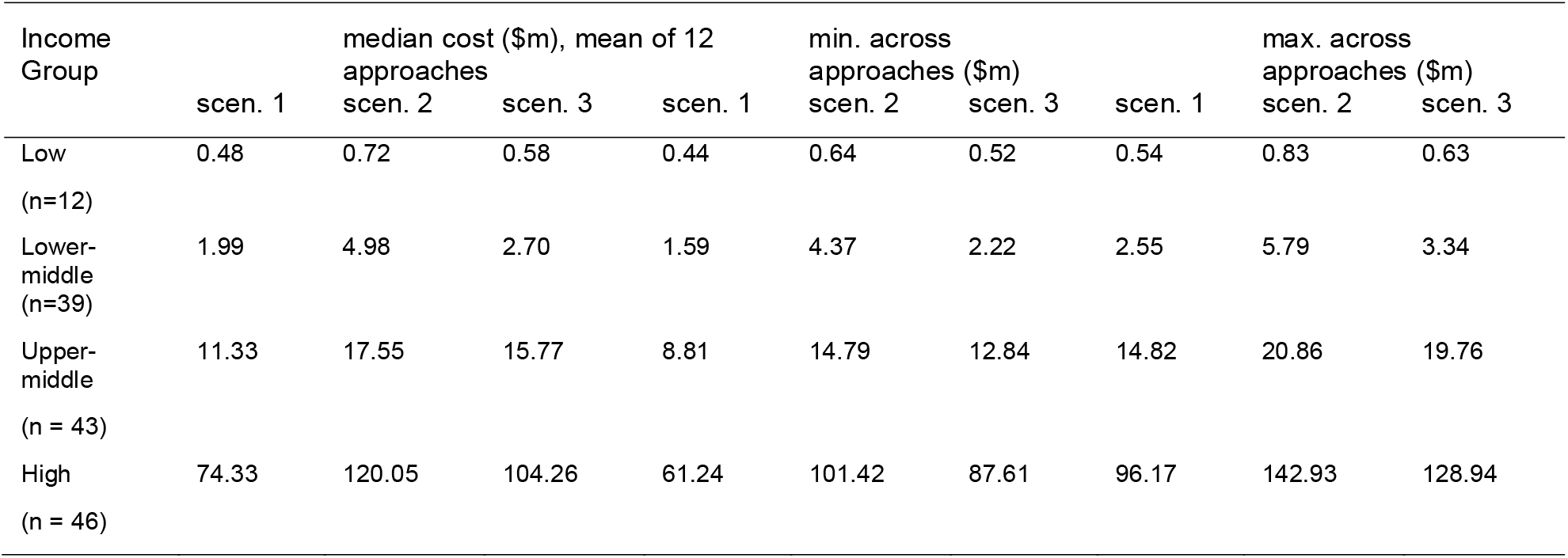
Median annual management cost per country of expanding to 30%, aggregated by World Bank income group. scen = scenario, $m = millions of U.S. dollars, min = minimum value for the lowest country in the group, max = maximum value for the highest country in the group

